# A collection of yeast cellular electron cryotomography data

**DOI:** 10.1101/594432

**Authors:** Lu Gan, Cai Tong Ng, Chen Chen, Shujun Cai

**Affiliations:** Department of Biological Sciences and Centre for BioImaging Sciences, National University of Singapore, Singapore 117543

**Keywords:** yeast, chromatin, nucleus, cryo-ET, cryo-EM, template matching, mining

## Abstract

**Background:** Cells are powered by a large set of macromolecular complexes, which work together in a crowded environment. The *in situ* mechanisms of these complexes are unclear because their 3-D distribution, organization, and interactions are largely unknown. Electron cryotomography (cryo-ET) is a key tool to address these knowledge gaps because it produces cryotomograms -- 3-D images that reveal biological structure at approximately 4-nm resolution. Cryo-ET does not involve any fixation, dehydration, staining, or plastic embedment, meaning that cellular features are visualized in a life-like, frozen-hydrated state. To study chromatin and mitotic machinery *in situ*, we have subjected yeast cells to a variety of genetic and/or chemical perturbations, cryosectioned them, and then imaged the cells by cryo-ET.

**Findings:** Every study from our group has generated more cryo-ET data than needed. Only the small subset of data that contributed to figures in these studies have been publicly shared. Here we share more than 1,000 cryo-ET raw datasets of cryosectioned budding yeast *S. cerevisiae*. This data will be valuable to cell biologists who are interested in the nanoscale organization of yeasts and of eukaryotic cells in general. To facilitate access, all the unpublished tilt series and a subset of corresponding cryotomograms have been deposited in the EMPIAR resource for the cell-biology community to use freely. To improve tilt series discoverability, we have uploaded metadata and preliminary notes to publicly accessible google spreadsheets.

**Conclusions:** Cellular cryo-ET data can be mined to obtain new cell-biological, structural, and 3-D statistical insights *in situ*. Because these data capture cells in a life-like state, they contain some structures that are either absent or not visible in traditional EM data. Template matching and subtomogram averaging of known macromolecular complexes can reveal their 3-D distributions and low-resolution structures. Furthermore, these data can serve as testbeds for high-throughput image-analysis pipelines, as training sets for feature-recognition software, for feasibility analysis when planning new structural cell-biology projects, and as practice data for students who are learning cellular cryo-ET.

## DATA DESCRIPTION

### Background

Cryo-ET is the combination of electron cryomicroscopy (cryo-EM) and computed tomography. In a cryo-ET experiment, 2-D cryo-EM data are incrementally recorded while the sample is rotated by typical angular steps of 1° to 3° over a range of -60° to +60°. These “tilt series” images are then mutually aligned and combined to generate a 3-D reconstruction called a cryotomogram. Because the cryotomogram contains a single field of view, cryo-ET is particularly valuable for the structural analysis of “unique” objects that cannot be averaged, such as cells [1-3]. A cryotomogram can contain a piece of tissue, cell, a portion of a cell, an isolated organelle, or a field of purified macromolecular complexes. This data note focuses on cryo-ET data of cryosectioned cells.

Cryo-EM is becoming a “big data” method [4]. Highly automated cryo transmission electron microscopes, automated data-collection software, and fast-readout direct-detection cameras can now generate terabytes of data per day [5-11]. Cryo-EM “single-particle analysis” (SPA) raw data contain many copies of conformationally and constitutionally similar macromolecular complexes that are suspended in buffer. In contrast, cellular cryo-ET raw data contain many species of macromolecular complexes. Furthermore, cellular cryo-ET data are usually recorded at lower magnification than for SPA. This dichotomy reflects (with exceptions) different goals: SPA studies aim to achieve maximum resolution of a few species of macromolecular complexes while cellular cryo-ET studies aim to determine how macromolecular complexes are distributed and organized in their intracellular environment. SPA and cellular cryo-ET studies do share similarities. Notably, only a small percentage of the collected data contribute to published models.

Our group has collected hundreds of tilt series per project. Because our studies are focused on one or a few types of structures, most of our data is in surplus. Two types of surplus data are “byproducts”, i.e., imaged cell positions that lack the targeted structures, and “bystanders”, i.e., imaged cellular structures adjacent to the targeted structures. We have previously shared cryo-ET data with collaborators and colleagues using commercial internet solutions like Dropbox and Google drive, but we found that these tools were suboptimal for sharing multi-gigabyte files. Alternative web technologies have allowed resources such as Electron Microscopy Public Image Archive (EMPIAR) [12] and the Caltech Electron Tomography Database (ETDB-Caltech) [13, 14] to share terabyte-sized datasets globally and more conveniently. We have deposited our published and surplus cryo-ET tilt series data in EMPIAR.

### Context

We are interested in the relationship between macromolecular structure and function inside cell nuclei. As a model system, we use yeast cells that are arrested at well-defined points in the cell cycle (Fig. 1A). We have shown that chromatin is packed irregularly without forming any monolithic condensed structures in both interphase and mitosis [15, 16] and that the majority of outer-kinetochore Dam1C/DASH complexes assemble as partial rings and do not contact the kinetochore microtubules’ curved tips *in situ* [17]. These studies show that the intracellular distribution and organization of macromolecular complexes are not always consistent with the models derived from *in vitro* studies. Indeed, our efforts to locate Dam1C/DASH *in situ* were hampered because we originally searched for complete rings butted up against curved microtubule protofilaments. We also had difficulty locating condensed chromosomes in fission yeast because we were expecting to find a monolithic nucleosome aggregates separated from a relatively “empty” nucleoplasm.

**Figure 1:**
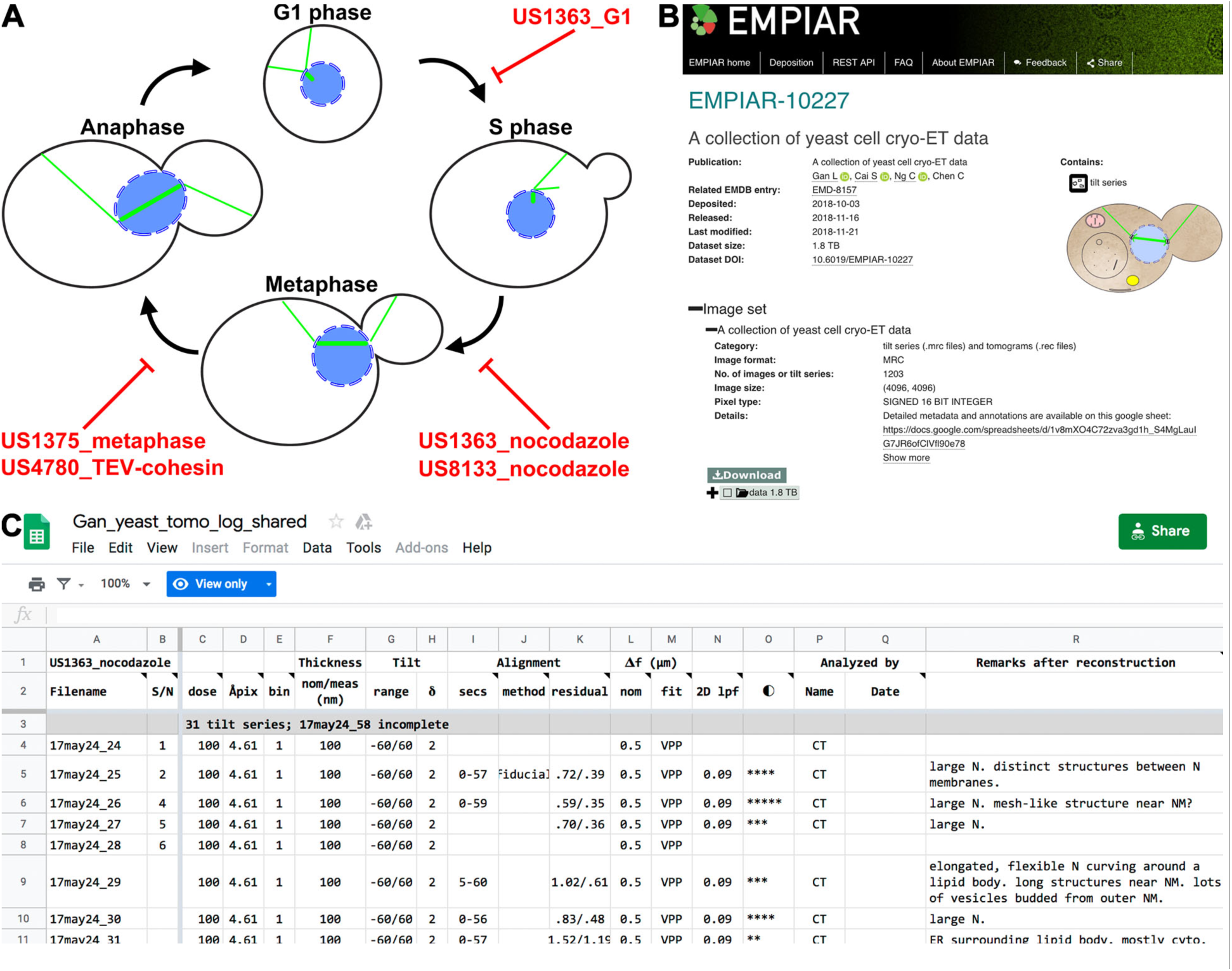
Yeast cryo-ET dataset summary. **(A)** Yeast cell-cycle stages sampled by this data. The red text indicates the strain ID plus either the cell-cycle state or treatment. **(B)** Screenshot of the EMPIAR entry. Downloads are faster and more reliable when done with the recommended client (Aspera Connect, as of this writing). **(C)** Screenshot of the preliminary notes, which are shared as google sheets named after the red text in panel A.

Our group has recorded more than one thousand tilt series of cryosectioned yeast cells. These include the budding yeast *Saccharomyces cerevisiae* and the fission yeast *Schizosaccharomyces pombe*. Only a minority of our recorded tilt series were presented in a paper; this data subset is already available at EMPIAR. Here we present the surplus cellular tilt series data we collected as part of published studies. We have neither analyzed nor intend to analyze in detail the vast majority of this data. These data will be valuable to other groups interested in macromolecular complexes and cytological features both within and outside the nucleus. Because the typical cryotomogram has ∼ 4-nm resolution, many structures can be identified on the basis of their shape, size, and intracellular context. The cryotomographic densities of some of these structures may contain features that are difficult to see in EM images of plastic sections. Notable examples are nucleosomes and some of the smaller or thinner components of the chromosome segregation and cell-division machineries.

### Dataset format and logistics

All cryo-ET data files are saved in the MRC format [18] under the accession code EMPIAR-10227 (Fig. 1B). Each dataset has a unique name, based on the date of data collection plus a serial number. For example, 18jun04a 02 is the second tilt series collected on 2018, June 4, session “a”. Future depositions may use the alternative YYYYMMDD_SN format, making the previous example 20180604_02. Some datasets include both a tilt series and a cryotomogram. The filename extensions follow IMOD conventions: “.mrc” for tilt series and “.rec” for cryotomograms. To conserve storage space and speed up file transfers, the tilt series and cryotomograms have been compressed with lbzip2, and therefore have the “.bz2” extension. The current entry does not include any movie or electron-counted data. In the future, electron-counted raw data will be stored as LZW-compressed .tiff files.

The pixel sizes used in the present data range from 4.6 to 9.1 Ångstroms. Most of the data were recorded on direction-detection cameras that have ∼ 16 million pixels in a 4,096 x 4,096 pixel array. The typical field of view therefore ranges from ∼ 2 to 4 µm squared. Most tilt series consist of approximately 61 images because we typically use a ±60° tilt range and a 2° tilt increment. The pixel intensity values in most tilt series data are stored as 16-bit unsigned integers, so the typical tilt series is ∼ 2 gigabytes.

We have shared via google sheets a read-only set of tabbed spreadsheets that contain metadata and preliminary notes and observations (Fig. 1C, link in Availability section). These spreadsheets are “live” documents and will be updated as new datasets are deposited. The first spreadsheet tab has a summary of all the data, links to additional related resources, commonly used commands, and a link to an online feedback form. Subsequent tabs contain detailed information on each tilt series, grouped by a strain ID and a treatment condition. For example, the “US1363_nocodazole” spreadsheet describes cryo-ET data of US1363 cells that were treated with the tubulin-polymerization inhibitor nocodazole.

In the detailed metadata spreadsheets, each row corresponds to one tilt series. Some cells were imaged by serial cryo-ET and therefore have the sequence number of each contributing tilt series noted in the “S/N” column. The other columns organize the data-collection parameters, appraisal of image contrast, diagnostic remarks on the data-collection session and quality, and a guess about the cytological features and macromolecular complexes present in the imaged cell. The accuracy of some of our annotations of cytological features is limited by our current cryo-ET and cell-biology knowledge, but will improve with both experience and especially user feedback. We anticipate that cell biologists will use the sorting function to shortlist the tilt series most salient to their studies.

## Methods

Cells were either grown in conditions that arrest populations at defined stages of the cell cycle or treated with drugs to perturb their cytology and cell-cycle progress. Because of our interest in mitosis and chromosome condensation, the present data capture cells in G1 phase, metaphase, and in mitosis with disrupted mitotic spindles. Liquid-cultured cells were collected by either centrifugation or vacuum filtration. These cells were then either high-pressure frozen or self-pressurized frozen in the presence of the extracellular cryoprotectant dextran. The frozen-hydrated cell block was sectioned in a cryomicrotome, producing a ribbon of cryosections. This cryosection ribbon was attached to either a continuous- or holey-carbon EM grid, which had been pre-coated with 10-nm-diameter gold nanoparticles. The cell cryosections were then imaged on a Titan Krios equipped with a direct detector, with or without Volta phase contrast. Additional details can be found in our earlier papers [15-17] and in the online spreadsheets.

Cryotomogram reconstruction, visualization, and analysis were done on a modern workstation computer with popular open-source software (Table 1). Radiation damage causes some cryosection positions to undergo non-uniform distortions, meaning that alignments were done using only the fiducials proximal to the structure of interest. Most of the tilt series were aligned using 4 to 12 fiducials coincident with the nuclei. To improve the visualization of other features, users should redo the alignment using only the fiducial markers closer to their structures of interest. If local alignment is not desired, the tilt series can be semi-automatically aligned using fiducials spread throughout the field of view and then reconstructed using software like Etomo and Protomo [19, 20]. Such tomograms tend to have uniform resolution at all positions where the cryosection is in contact with the carbon substrate.

**Table 1:** Recommended hardware and software

| Tool | Recommendation | Notes |
| --- | --- | --- |
| Computer | Modern workstation | More memory (RAM) facilitates comparisons of multiple cryotomograms. |
| Display | 27+ inch monitor |  |
| Operating system | Linux | Most cryo-EM software is developed on Linux; extra effort is needed to run this software in Mac OS or Windows |
| Visualization software | 3dmod (IMOD) | FIJI can also be used, but it is not optimized for tomography data. |
| Reconstruction software | Etomo (IMOD) | A solid-state disk and a CUDA-compatible GPU are highly recommended |
| Download client | Aspera Connect | Fast, fault-tolerant software for large downloads from EMPIAR |
| Notes | Google sheets, Microsoft Excel | The shared spreadsheet can be downloaded and then customized. |

Cryotomograms are noisier than SPA reconstructions, meaning that these datasets are very difficult to comprehend when visualized as isosurfaces. Instead, cryotomograms are better visualized as tomographic slices: 2-D images that average multiple voxels along one axis. The slice thickness should match the structure of interest, e.g., 10 nm for nucleosomes. To facilitate comparison between multiple datasets, multiple cryotomograms can be simultaneously loaded into random-access memory in one instance of the program 3dmod [19]. Assuming they all “fit” into memory, cryotomograms loaded this way can be rapidly toggled in sequence using the “1” and “2” shortcut keys.

Reconstructed cryotomograms are usually the starting point of more quantitative analysis. Examples of deeper analysis by template matching, classification, and subtomogram averaging can be found in recent reviews and the many excellent papers cited within [21-24]. Because structural cell biology is a new field, most of our studies have required new analysis tools. We have written a number of python scripts to facilitate the 3-D packing analysis of subtomograms (https://github.com/anaphaze/ot-tools). These scripts control programs from mostly open-source image-analysis packages [19, 25-27].

### Data validation and quality control

The data have been recorded under a variety of conditions (magnification, dose, tilt increment, defocus) with different contrast mechanisms (defocus phase contrast vs. Volta phase contrast). Furthermore, the tilt series have differences in quality due to variations in either freezing, attachment to the grid, and radiation damage. Owing to this variability, we cannot assign a single validation metric to the entire set of tilt series. We have qualitatively assessed each tilt series’ contrast relative to others recorded in the same session (tens of tilt series per session). The contrast is rated from one to five stars and is recorded in the online spreadsheet columns marked with the “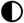 ” symbol. Four-to five-star data typically reveal features like membrane leaflets, clear separation of nucleosome-like particles, and particles smaller than nucleosomes. These evaluations were made either from tomograms when possible.

The deposited cryotomograms should be considered preliminary for three reasons. First, most of the cryotomograms were reconstructed using the subset of fiducial markers coincident with the nucleus, which results in lower reconstruction quality elsewhere in the cell. Second, the fiducial centers were manually fine-tuned for the few tilt series that contributed to the final published figures. Third, we anticipate that future developments in fiducial-assisted and fiducial-less alignment will produce better cryotomograms than currently possible.

### Re-use potential

The deposited yeast cryo-ET data contain a large number easy-to-find or abundant organelles and macromolecular complexes such as mitochondria, eisosomes, cytokinetic machinery, microtubule-organizing centers, fatty acid synthases, proteasomes, vacuoles, rough endoplasmic reticulum, lipid bodies, and cytoplasmic aggregates (Fig. 2). Cell biologists may want use these data to measure local concentrations of macromolecular complexes, detect interactions between these complexes, determine the orientations of large complexes *in situ*, test for the existence of putative cellular features, and determine how cellular bodies make direct contact with one another. Closer inspection may reveal poorly documented subcellular features. Examples of such features include mitochondrial filaments (Fig. 2A), ordered layers in lipid-droplet-like bodies (Fig. 2I), and amorphous cellular aggregates (Fig. 2J). Furthermore, this data will provide morphological, distance, or stoichiometric constraints for groups attempting to reconstitute either a complex or a cellular body.

**Figure 2:**
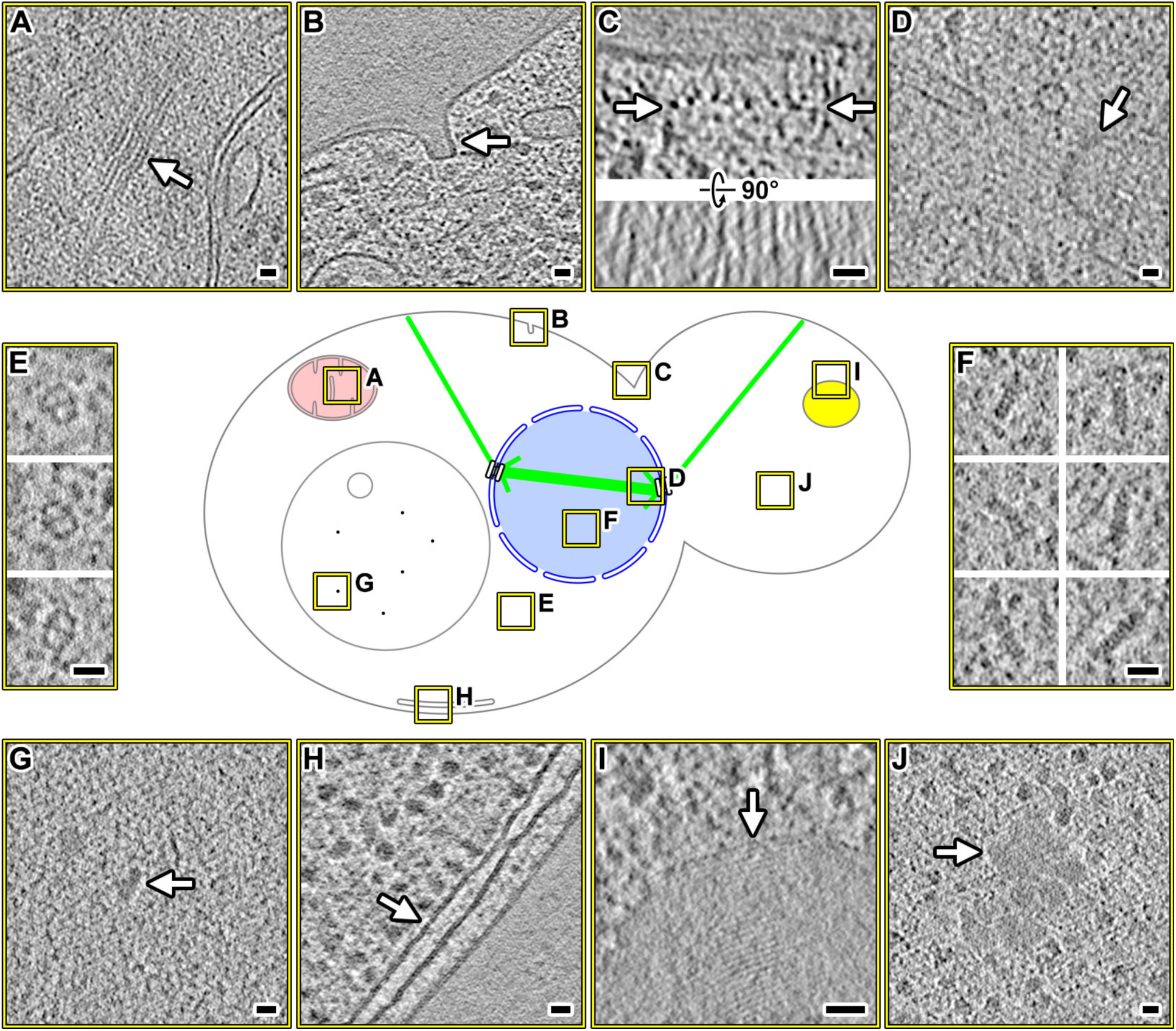
Easy-to-find structures in yeast cryotomograms. Center: graphical legend showing the locations of interesting features (boxed in yellow), which are enlarged as cryotomographic slices (10 - 20 nm thick). **(A)** Filament bundle within a mitochondrion. **(B)** Eisosome; see [30] for identification details. **(C)** Cytokinetic machinery. Upper panel: transverse view. The row of filamentous complexes is indicated by arrows. Lower panel: longitudinal view of the filaments. **(D)** Microtubule-organizing center. **(E)** Fatty acid synthases. **(F)** Intranuclear proteasomes. **(G)** Particles in a vacuole. **(H)** Endoplasmic reticulum adjacent to the plasma membrane. **(I)** Lipid-droplet-like body with periodic internal structure. **(J)** Amorphous cytoplasmic mass. Scale bar = 20 nm in all panels.

Higher-resolution structural information can be obtained by alignment and averaging of subtomograms containing copies of the macromolecular complex. If multiple copies of a macromolecular complex can be detected in one or more cellular cryotomograms, they can be analyzed as “single particles” and averaged together to achieve density maps that have high-resolution features, as discussed in recent reviews [21-24]. The centers of mass and orientation information can then be used to remap the average back into a volume the size of the cryotomogram. If the complexes are densely packed, these remapped models will reveal higher-order structure as seen in polysomes and oligonucleosomes [28, 29].

Cellular cryotomograms also contain hard-to-find structures (Fig. 3). These structures are either rare or they are located in cellular positions that we rarely target, such as the bud neck (Fig. 3C). Many of these structures, such as inter-membrane contact sites (Fig 3F) and lipid-body protrusions (Fig. 3H) are poorly documented in the cryo-ET literature. We anticipate that yeast cryo-ET data will help stimulate the discovery and detailed characterization of interesting eukaryotic subcellular bodies just as cellular cryo-ET has done for bacterial cell biology [31-35]. Furthermore, structures that are identified by other groups can be retrospectively analyzed in this data in the context of known cell-cycle states and pharmacological perturbations.

**Figure 3:**
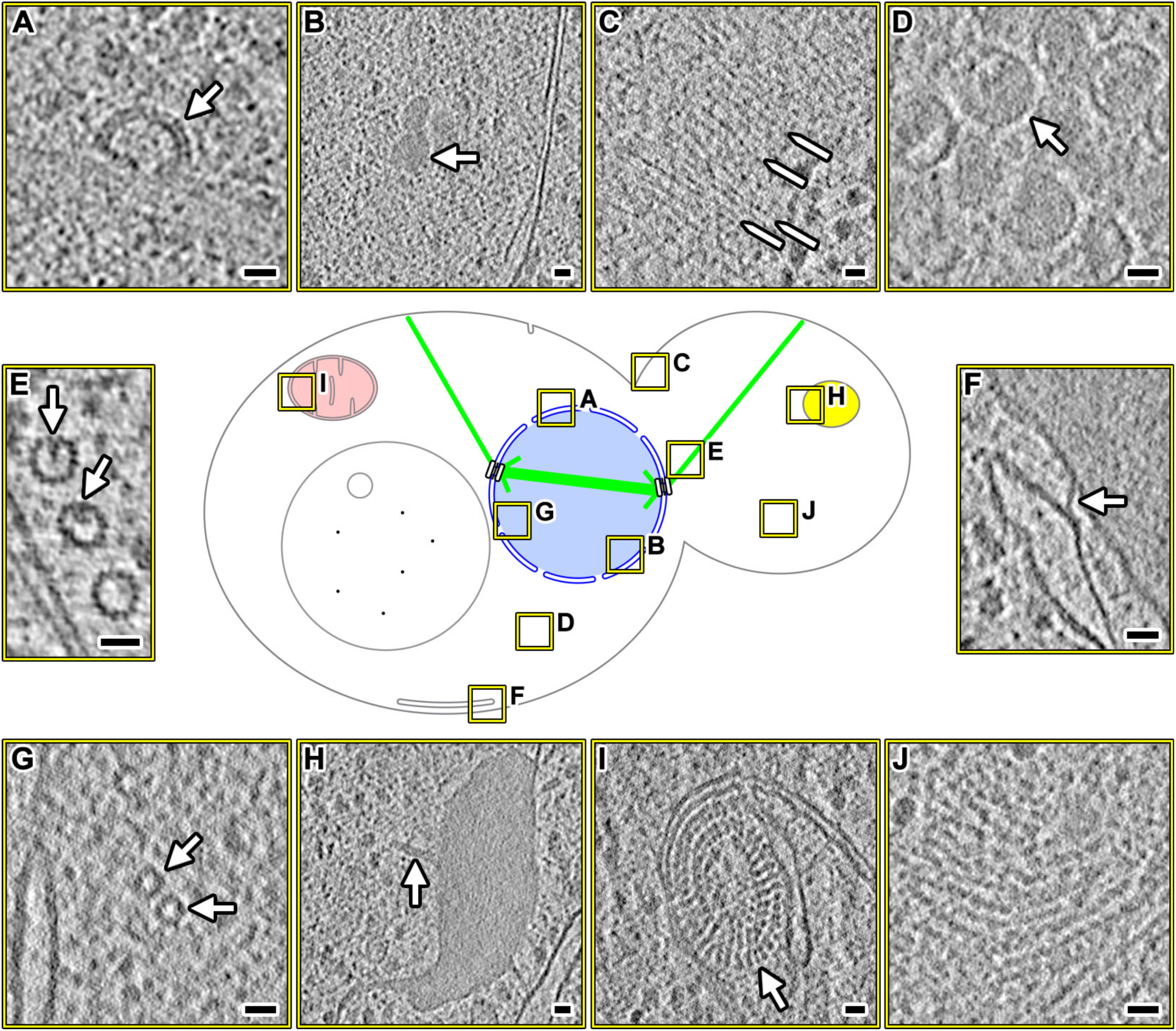
Hard-to-find structures in yeast cryotomograms. Center: graphical legend showing the locations of interesting features (boxed in yellow), which are enlarged as cryotomographic slices (10 - 20 nm thick). **(A)** A coated pit-like structure, docked to the outer nuclear membrane. **(B)** Intranuclear granule. **(C)** Septin-like cytokinesis machinery. A few examples are indicated by the pointed lines. These filaments run parallel to the mother-daughter cell axis. **(D)** Virus-like particles in the cytoplasm. **(E)** Luminal particles in cytoplasmic microtubules. **(F)** Connection between the endoplasmic reticulum and plasma membrane. **(G)** Short intranuclear 15-nm diameter tubes. **(H)** A lipid body with thin protrusions, one of which is indicated by the arrow. **(I)** Mitochondrial periodic structures extending from the inner membrane into the matrix. **(J)** Filamentous cytoplasmic aggregates.

Users should note we arrested the yeast cells in various cell-cycle stages to allow comparative studies of nuclear structures like chromatin, spindles, and kinetochores. Because the cell cycle affects the entire proteome, this dataset will therefore shed light on how other organelles and cytoplasmic macromolecular complexes are cell-cycle regulated. Some of the structures observed in this yeast data may also be stress-induced. Indeed, recent studies showed that upon starvation, eukaryotic translation initiation factor 2B forms large filament bundles in budding yeast [36, 37].

These data span a range of defoci and magnifications, with or without the Volta phase contrast [38]. Such experimental diversity will allow software developers to test the robustness of new image-processing routines used in automated alignment [19, 20], template matching (also called 3-D particle picking), subtomogram averaging and classification [27, 39-42]. The yeast cryo-ET data can also be used to train machine-learning algorithms to detect features in both tilt series and cryotomograms [43-45]. Furthermore, data-sharing resources may use this data to develop annotation and browsing tools [46].

The vast majority of our cryo-ET imaging was recorded with first generation direction-detection cameras, without energy filtering. If either the structure of interest or a structure of equivalent size can be detected in the present data, then it will most certainly be detectable in data recorded on electron-counting cameras, both with or without energy filtering. Therefore, these data will facilitate feasibility analyses.

Finally, new structural cell biologists will find these data useful as real-world examples that complement the lessons from cryo-EM tutorials [47, 48]. The vast majority of the deposited data are from grids that have gold nanoparticles, making the alignment process similar to -- and therefore a direct follow-on to the IMOD plastic-section tutorial dataset [48]. Students can use the reconstructed tomograms to practice manual annotation and more automated analyses such as template matching and subtomogram averaging.

### Availability of supporting data

We have deposited this data under accession code EMPIAR-10227 (https://dx.doi.org/10.6019/EMPIAR-10227). We excluded “unusable” tilt series, which have the following image or sample properties: extreme drift, occlusion by large ice crystals, cracks in the ice or carbon substrate, or completely detached sections. We also included a copy of the tilt series that were already deposited as part of original research papers. Key metadata are available in read-only google sheets (https://goo.gl/mwWyTk), which can be copied to the user’s own google drive or downloaded as a spreadsheet file. Thereafter, the user can sort the rows to identify smaller subsets of tilt series that have the desired properties or structures. Feedback can be sent via a google form (https://goo.gl/forms/FtU8RtbXCfbAa2gn2).

The tilt series and cryotomograms are organized in the following directory structure:

~~~
Sample_ID_1
 Session_ID
  Tilt_series
   series_first.mrc.bz2
   …
   series_last.mrc.bz2
  Tomograms
   series_first.rec.bz2
   …
   series_last.rec.bz2
Sample_ID_2
 Session_ID
  Tilt_series
    series_first.mrc.bz2
   …
   series_last.mrc.bz2
  Tomograms
   series_first.rec.bz2
   …
   series_last.rec.bz2
~~~

## Abbreviations

cryo-EM: cryo-electron microscopy / electron cryomicroscopy
cryo-ET: cryo-electron tomography / electron cryotomography

## Competing interests

The authors do not have any competing interests.

## Funding

Singapore Ministry of Education T1 R-154-000-A49-114 and T1 R-154-000-B42-114.

## Authors’ contributions

Experiments: CTN, CC, SC. Metadata organization and writing: LG.

## Acknowledgements

We thank Gemma An and Chithran VM for help with some reconstructions; Uttam Surana and Mohan Balasubramanian for the yeast strains; Ardan Patwardhan and Andrii Iudin for feedback on data organization; Christoph Baranec for discussion on astronomy data-sharing practices; Paul Matsudaira, Jian Shi, Ann Tran, and Ping Lee Chong for setting up and operating the cryo-EM platform at the National University of Singapore Centre for BioImaging Sciences; and our many colleagues for discussions on interesting cell-biology questions.

